# AI-assisted protein design to rapidly convert antibody sequences to intrabodies targeting diverse peptides and histone modifications

**DOI:** 10.1101/2025.02.06.636921

**Authors:** Gabriel Galindo, Daiki Maejima, Jacob DeRoo, Scott R. Burlingham, Gretchen Fixen, Tatsuya Morisaki, Hallie P. Febvre, Ryan Hasbrook, Ning Zhao, Soham Ghosh, E. Handly Mayton, Christopher D. Snow, Brian J. Geiss, Yasuyuki Ohkawa, Yuko Sato, Hiroshi Kimura, Timothy J. Stasevich

## Abstract

Intrabodies are engineered antibodies that function inside living cells, enabling therapeutic, diagnostic, and imaging applications. While powerful, their development has been hindered by challenges associated with their folding, solubility, and stability in the reduced intracellular environment. Here, we present an AI-driven pipeline integrating AlphaFold2, ProteinMPNN, and live-cell screening to optimize antibody framework regions while preserving epitope-binding complementarity-determining regions. Using this approach, we successfully converted 19 out of 26 antibody sequences into functional single-chain variable fragment (scFv) intrabodies, including a panel targeting diverse histone modifications for real-time imaging of chromatin dynamics and gene regulation. Notably, 18 of these 19 sequences had failed to convert using the standard approach, demonstrating the unique effectiveness of our method. As antibody sequence databases expand, our method will accelerate intrabody design, making their development easier, more cost-effective, and broadly accessible for biological research.

## INTRODUCTION

Intrabodies are a special class of antibodies that are engineered to fold and function within the intracellular environment^1,2^. This property gives intrabodies great therapeutic and diagnostic potential because they can bind target biomolecules deep within living cells and tissue. Furthermore, when intrabodies are tagged with fluorescent proteins, they become powerful imaging reagents that can bind and light up challenging targets that cannot be imaged with standard genetic fusion tags like GFP. For example, fluorescent intrabodies have been used to image conformational^3–6^ and linear epitopes^7,8^, macromolecular complexes^9,10^, and chemical modifications^11–13^.

Despite the broad applicability of intrabodies, engineering them has proven to be a significant challenge. The antibody scaffolds from which intrabodies are derived have evolved to function in extracellular environments like the bloodstream, where conditions support the formation of disulfide bonds that are critical for maintaining structure. In contrast, the reduced intracellular environment disrupts these bonds, often leading to issues such as misfolding, aggregation, and degradation^14–16^. Additionally, the intracellular environment features numerous differences from the extracellular environment in terms of off-target binding and aggregation. For these reasons, the straightforward placement of antibody domains into an intrabody format – whether it be a single-chain variable fragment^17^ (scFv) or a nanobody^2,18–20^ – often results in a non-functional protein. Thus, despite the wide availability of antibodies commercially and in protein databases, there are relatively few functional intrabodies^21,22^.

One specific area of research that could benefit from additional intrabodies is the study of histone modifications^11^. These reversible chemical marks – which include acetylation, methylation, and phosphorylation, among others – decorate the unstructured tails of nucleosomes across the genome. The modifications are believed to work both individually and in combination to help shape chromatin architecture and regulate gene expression^23,24^. As such, they are strongly correlated with human health and disease, making them promising therapeutic targets^25,26^. However, because the marks are reversible, they are difficult to image in living cells using standard genetic tags^27,28^. Instead, specialized imaging probes like intrabodies are needed that can dynamically bind and light up the modifications as they come and go. Although there has been some progress developing such probes^27,28^, the pace has been slow due to the inherent engineering challenges. For this reason, the dynamics of histone modifications have remained elusive, as has their precise roles in gene regulatory networks.

Recent advances in artificial intelligence, including tools like AlphaFold2^29^, RFdiffusion^30^, and protein large language models^31^, offer new opportunities to overcome these challenges. These technologies not only enable the *de novo* creation of proteins with novel functions^32–34^, but also provide a powerful new platform to improve existing proteins like antibodies. While numerous efforts have focused on optimizing the complementarity determining regions of antibodies to improve target binding affinity^35–37^ or the optimization of therapeutic antibody sequences to minimize self association and off-target binding under standard formulation conditions^38,39^, it remains unclear if these technologies can similarly enhance the framework regions of antibodies to improve their functionality when expressed as intrabodies in living cells.

To address this issue, we here develop an AI-driven approach to accelerate the development of functional intrabodies. Using a novel pipeline that leverages antibody domain annotation^40^, AlphaFold2^29,41^, ProteinMPNN^42^, and live-cell screening, we demonstrate how existing antibody sequences can be rapidly converted into functional intrabodies. As a proof-of-principle, we first develop two peptide-binding intrabodies, one targeting the synthetic FLAG tag^43^ and the other targeting a linear epitope of the SARS-CoV-2 nucleocapsid protein^44^. We then turn our attention to endogenous histone modifications. To highlight the speed and practicality of our approach, we create a panel of intrabodies that bind diverse residue-specific chemical modifications on both histones H3 and H4, including H3T3ph, H3K4me1, H3K4me2, H3K4me3, H3K9(un), H3S10ph, H3K14ac, H3K27me1, H3K27me2, H3K27ac, H3K36me3, H4K5ac, H4K8ac, H4K12ac, and H4K16ac, tripling the number of intrabodies available for this purpose^27^. We demonstrate the functionality of these intrabodies in living cells and provide their sequences so they can be further refined or modified to image or manipulate histone modifications *in vivo*. Furthermore, we provide open-source code implementing our pipeline, and provide useful metrics and tables to help predict which candidate designs will retain functionality inside cells. As the number of published antibody sequences continues to rapidly grow, we believe our approach will significantly increase the number, utility, and accessibility of intrabodies to scientists in the life sciences.

## RESULTS

### Developing a pipeline for transforming existing antibody sequences into enhanced scFvs for intracellular applications

We developed a pipeline to explore the potential of AI-based protein design tools to enhance scFv engineering (Fig. 1A). We were driven by failures to produce a functional scFv based on the commercial M2 anti-FLAG antibody, whose peptide sequence had recently been published^45^. Initially, we reformatted the M2 sequence into a standard scFv with a flexible glycine-serine-rich linker. However, when fused to GFP and co-expressed with FLAG-tagged histone H2B, the scFv failed to colocalize with its target, instead showing uniform cellular distribution – a clear indication of it remaining unbound and freely diffusing (Fig. 1B). We attempted to fix this problem by loop grafting the M2 complementary determining regions (CDRs) onto a stable scFv framework region that we had previously used to generate anti-HA^7^ and anti-FLAG^46^ ‘frankenbodies’. We also tested a new scFv framework region derived from an anti-Pol II scFv^47^, selected due to its high sequence similarity to the M2 frameworks. Unfortunately, none of these variants achieved specific colocalization with FLAG-tagged H2B (Fig. S1A).

**Fig 1.**
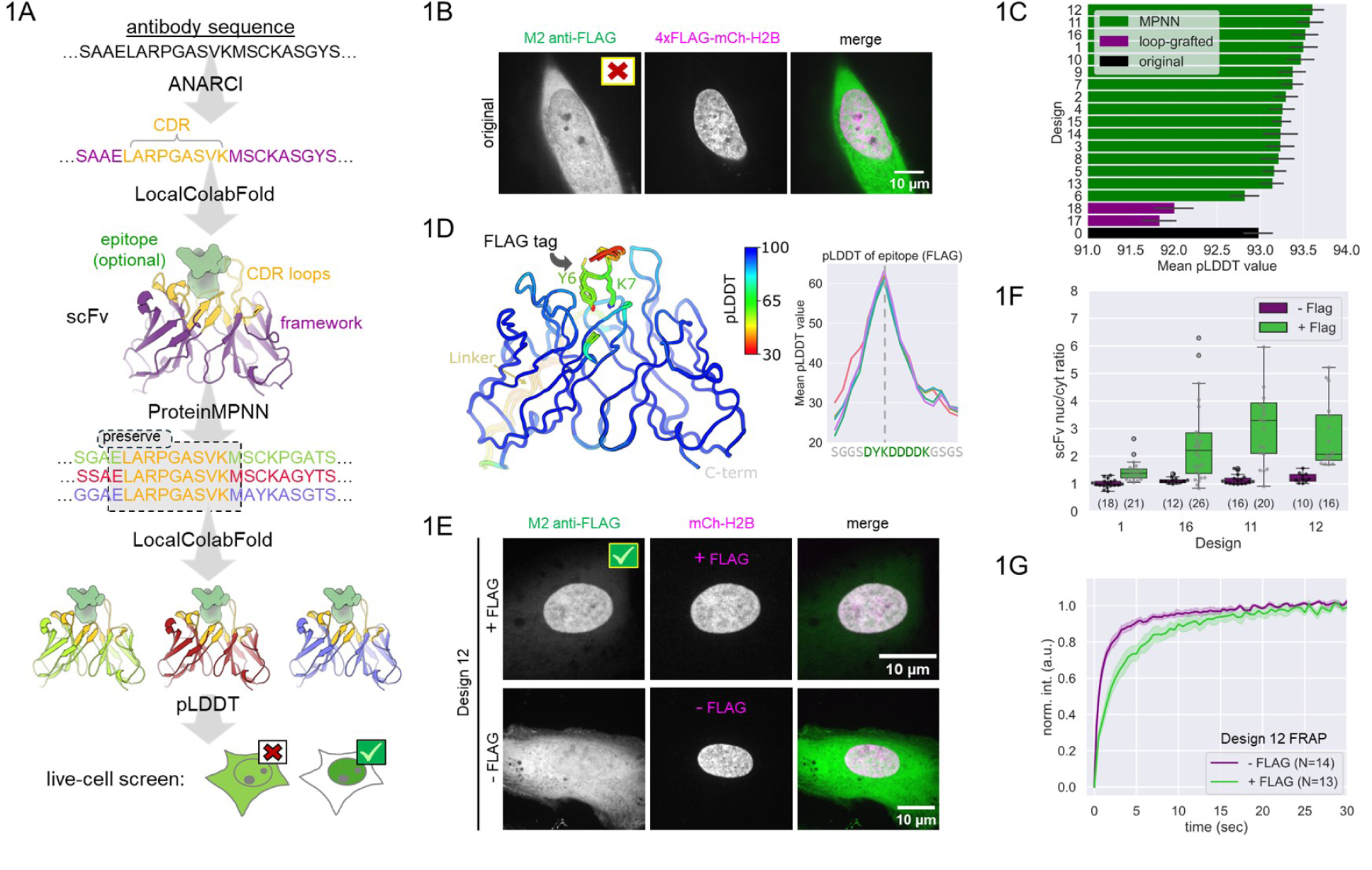
Pipeline to convert antibody sequences into functional intrabodies. **A.** Schematic showing the pipeline, from antibody sequence to live-cell screening. **B.** The original M2 anti-FLAG scFv (green) does not colocalize with 4xFLAG-mCh-H2B (magenta) in the nucleus. **C.** The ranked pLDDT scores of all M2 anti-FLAG scFv designs (includes scFv and FLAG epitope), including the original (black bar), two loop-grafted variants (purple), and 16 from the pipeline (green). Error bars show the standard deviation from LocalColabFold predicted models (N=5 each). **D.** The top four predicted structures (models 1, 16, 11, and 12) overlaid with the original M2 anti-FLAG structure (model 0), colored by pLDDT. Right, the corresponding overlaid pLDDTs of the tested binder, with the FLAG tag (DYKDDDDK) indicated in green text. **E.** Sample cell from the live-cell screen of design 12. This scFv (green) colocalizes with the FLAG-tagged H2B (magenta). When there is no FLAG-tagged protein co-expressed, the scFv shows uniform localization (bottom). **F.** Live-cell measurement of the nuclear to cytoplasmic ratio of the indicated scFv designs (number of cells (N) indicated; box is 25-75% interquartile range; whiskers mark points within 1.5✕ the interquartile range). **G.** FRAP experiments in the nucleus of cells expressing design 12 scFv in the presence and absence of FLAG-H2B. The number of cells (N) is indicated; the shaded region shows the 95% CI.

Given the widespread use of the M2 antibody, we were convinced our scFv failed in living cells due to misfolding or insolubility, rather than a failure in antigen binding and specificity. We hypothesized that if we could design more soluble scFv framework regions based on the original sequence, then this would enhance their compatibility with the original CDRs, thereby increasing the likelihood of successful loop grafting. To test this hypothesis, we created a computational pipeline that integrates several AI-based tools. The pipeline begins by annotating the antibody sequence to extract the CDRs and framework regions using ANARCI^40^. Second, we predict the 3D structure of the scFv:target complex using LocalColabFold^41^, which combines AlphaFold2^29^ with the fast multiple sequence generation method MMseqs2^48,49^. Third, we input the predicted structure into ProteinMPNN^42^, holding regions near the annotated CDRs fixed. For the remaining residues, ProteinMPNN attempts to predict favorable new amino acid selections. Crucially, the resulting ProteinMPNN framework sequence differs from the original, but tends to encode a protein that is more well behaved because the underlying neural network has been trained on well-behaved, soluble proteins with solved crystal structures^42,50^. We hypothesized that these new framework sequences would be preferable with respect to combating off-target aggregation of the designed proteins, while maintaining the critical 3D structure needed for compatibility with the original CDRs.

Running the M2 anti-FLAG scFv through the pipeline resulted in 15 designs. Compared to the original M2 anti-FLAG scFv and the two unsuccessful loop-grafted versions, the new designs exhibited considerably improved pLDDT (predicted local distance difference) scores (Fig. 1C). AlphaFold2 predicted 3D structures of these designs closely matched the original model (Fig. 1D, left). While the per-residue confidence (pLDDT) was significantly lower for the FLAG epitope than the scFv (Fig. 1D) (excluding the linker), we noted that the confidence in the authentic epitope was nonetheless higher than the confidence for flanking residues, hinting at latent detection of binding specificity by AlphaFold2. Notably, the pLDDT scores for the FLAG epitope were significantly higher than those for adjacent residues (Fig. 1D, right), even when the FLAG epitope was challenged with a nearby HA epitope (Fig. S1B), indicating binding specificity. Furthermore, the predicted orientation of the FLAG epitope matched that of a recently determined structure based on the full-length parental M2 antibody (8RMO)^51^, despite being predicted prior to the deposition of the structure in the Protein Data Bank (Fig. S1C).

We next screened the top four designs in living cells, selected based on pLDDT scores (designs 12, 11, 16, and 1 in Fig. 1C). Remarkably, all four candidates successfully colocalized with FLAG-H2B in the nucleus, as quantified by the ratio of nuclear to cytoplasmic fluorescence (Fig. 1E,F). The specificity of the best scFv (design 12) was further validated in cells without FLAG tags, where the scFv displayed homogenous localization, indicating minimal off-target binding (Fig. 1E, bottom; Fig. 1F, purple). Consistently, FRAP on design 12 was significantly slower in the presence of FLAG-H2B (Fig. 1G), with a recovery half-time of ∼2 seconds. The timescale of this recovery is similar to FRAPs we performed on monovalent antibodies (Fab) generated from full-length antibodies with affinities in the 10 nM range^52^. Consistent with this comparison, the affinity of the full-length M2 anti-FLAG antibody was recently estimated to be ∼2 nM^51^. These data therefore suggest the CDRs of our M2 scFvs remain intact despite the large number of changes we made to their framework regions. This underscores the success of our pipeline in generating functionally improved scFvs that maintain specific antigen recognition.

### Using the pipeline to create an intrabody against the SARS-CoV-2 nucleocapsid protein

To determine if our technique could be generalized, we next targeted a medically relevant antigen: the nucleocapsid protein (NP) of SARS-CoV-2. The NP is a promising diagnostic target due to its early and abundant expression during viral infection^53^. We had previously developed and sequenced an anti-nucleocapsid mouse monoclonal antibody (mBG17), identifying its epitope as a short 10-amino acid peptide (EpNP: DDFSKQLQQS) that forms an alpha-helical structure^44,54^. Similar to our experiences with the M2 anti-FLAG scFv, when we attempted to convert this sequence into an intracellularly functional scFv or frankenbody, we were unsuccessful (Fig. S2A). These previous failures made the anti-EpNP scFv an ideal candidate to further test our computational pipeline. Furthermore, unlike the FLAG epitope and M2 anti-FLAG antibody, which are well-characterized and supported by crystal structures available in the Protein Data Bank, the mBG17 antibody lacks structural information, offering an unbiased test case for evaluating the predictions of our pipeline.

Mirroring our work with the M2 anti-FLAG antibody sequence, we generated 45 new scFv sequences targeting the NP epitope. All had higher pLDDT scores than the original, with some designs better than others (Fig. S2B). To assess how predictive these pLDDT scores were for live-cell functionality, we focused on six designs across the pLDDT score range (Fig. 2B). The predicted 3D structures for these six designs showed consistent placement of the epitope within the CDR binding pocket (Fig. 2C), with the central portion having especially high pLDDT values (Fig. 2C). This aligned well with our earlier work^44^ that identified the key residues within the NP epitope that are critical for binding to the antibody, bolstering our confidence in the predicted structures.

**Fig. 2.**
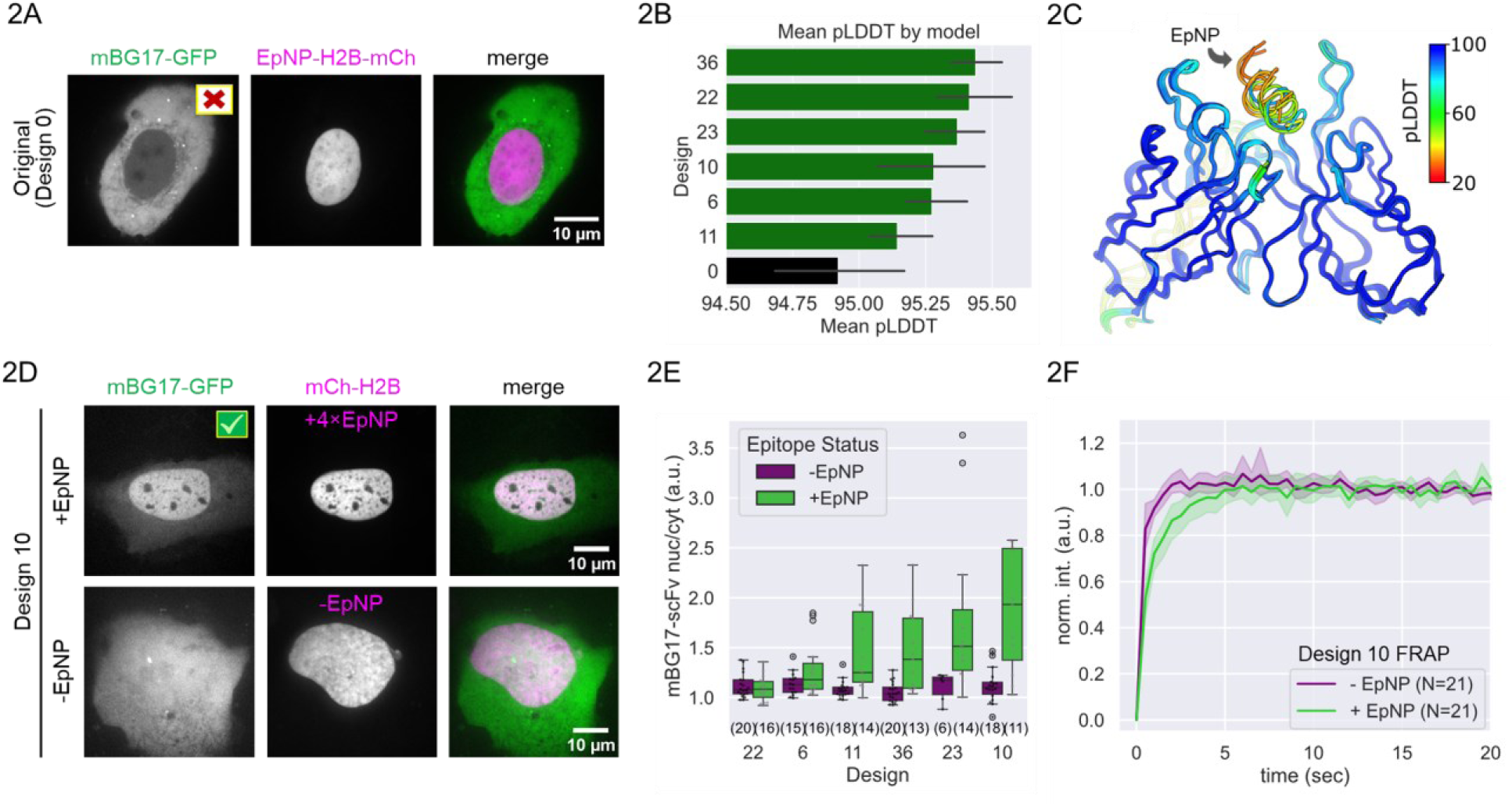
Engineering a functional intrabody against the nucleocapsid protein from SARS-CoV-2. **A.** An scFv (mBG17-GFP; green) based on the sequence of a wildtype antibody against the SARS-CoV-2 nucleocapsid protein does not colocalize with its target epitope (EpNP-H2B-mCh, magenta) in living cells. **B.** The ranked pLDDT scores of a selection of mBG17 pipeline designs (includes both scFv and EpNP; green bars) along with the original (black bar). Error bars show the standard deviation from LocalColabFold predicted models (N=5 each). **C.** The predicted structures (models 11, 6, 10, 23, 22, and 36) overlaid, colored by pLDDT. **D.** Design 10 (green) colocalizes with its target epitope (top; +EpNP = EpNP-mCh-H2B; magenta) and displays uniform fluorescence throughout cells in the absence of its target epitope (bottom; -EpNP = mCh-HB; magenta). **E.** Live-cell measurement of the nuclear to cytoplasmic ratio of the indicated scFv designs (number of cells (N) indicated; box is 25-75% interquartile range; whiskers mark points within 1.5✕ the interquartile range). **F.** FRAP experiments in the nucleus of cells expressing design 10 mBG17 scFv in the presence (+EpNP = EpNP-mCh-H2B) and absence (-EpNP = mCh-H2B) of EpNP. The number of cells (N) is indicated; the shaded region shows the 95% CI.

To screen individual designs, we co-transfected each in cells together with a plasmid encoding a 4×EpNP-tagged histone H2B. Three of the six tested anti-EpNP scFv designs successfully colocalized with the EpNP-H2B in the nucleus (Figs. 2D,E). Interestingly, the successful candidates were not those with the highest pLDDT scores, suggesting that high scores alone might not always predict solubility and functional success. To verify specific binding, we expressed the successful scFv in cells lacking H2B-EpNP. In these cells, we observed a uniform distribution, indicating that the scFv was freely diffusing and unbound, with no detectable off-target binding (Fig. 2D, lower and Fig. 2E, purple). Fluorescence recovery after photobleaching (FRAP) further confirmed this specificity, showing significantly slower recovery rates in cells containing H2B-EpNP compared to those without it (Fig. 2F).

In our analysis of the sequence alignment of the screened scFv variants, we identified a feature that correlated with success: residue 80 was K in more effective designs versus S or N in less effective designs (Fig. S2C,D). This residue is situated in a beta hairpin turn outside the CDR loops, and could play a role in stabilizing CDR conformations (Fig. S2D). Intriguingly, our previous studies that utilized random mutagenesis on different scFvs also identified residue 80 as crucial for scFv solubility^55^. However, the presence of K80 alone is not sufficient for enhanced performance, as the original sequence also contained K80. This suggests that the improved solubility and functionality likely result from modifications to multiple residues. Nonetheless, these observations highlight the effectiveness of our strategy in rapidly generating functional scFv sequences and also demonstrates how a comparison of different designs can be used to quickly generate hypotheses about key residues.

### Developing intrabodies targeting post-translational histone H3 modifications

Encouraged by these successes, we next set out to develop intrabodies that target endogenous post-translational histone modifications. We refer to such modification-specific intracellular antibodies as ‘mintbodies’^13,27^. Mintbodies dynamically bind their target modifications, enabling real-time imaging and tracking in living cells without affecting cell divisions and animal development^27,56^.

Since 2013, five mintbodies against histone and RNA polymerase II modifications have been developed^13,47,55,57,58^. The first two were selected from several hybridoma clones^13,57^ and the other three involved random and/or site-directed mutagenesis with extensive screening^47,55,58^. In the latter cases, the original scFv were only a few mutations away from being functional in the intracellular environment, suggesting they may have been fortuitously close to working from the start. Based on our experience with anti-EpNP, we suspect other scFvs may require many mutations before they can properly fold and function inside cells, in which case random mutagenesis becomes impractical due to the immense combinatorial possibilities. We therefore hoped our computational pipeline could significantly reduce the search space and expedite the production of useful mintbodies.

To test this, we decided to focus on a set of mouse monoclonal antibodies targeting three distinct modifications to histone H3: Lysine 27 acetylation (anti-H3K27ac, clone 9E2H10/CMA309)^59^, Serine 10 phosphorylation (anti-H3S10ph; clone 7G1G7/CMA311)^60^, and Lysine 9 trimethylation (anti-H3K9me3; clone 2F3/CMA318)^61^. These antibodies are excellent candidates for our pipeline because – despite being sequenced and well characterized – each fails to bind its target modification in living cells when converted into an scFv format (Fig. 3A). Furthermore, the structures of these antibodies have not been experimentally determined, so our pipeline predictions should be free from any bias that could arise from AlphaFold having prior access to the data during its training.

**Fig. 3.**
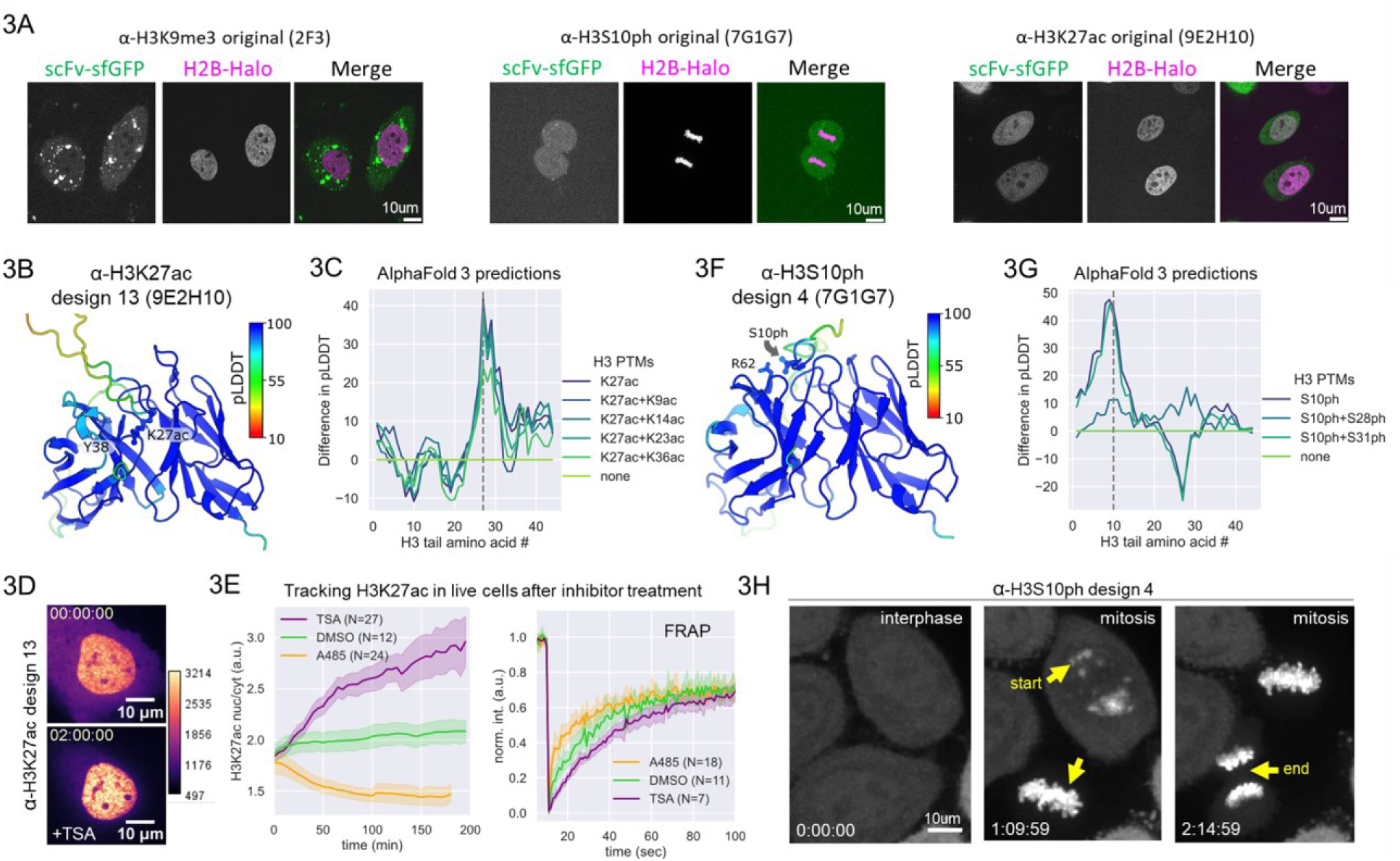
Intrabodies targeting post-translational histone H3 modifications. **A.** Three original scFvs (green) against histone H3 Lysine 9 trimethylation (H3K9me3; 2F3), Serine 10 phosphorylation (H3S10ph; 7G1G7), and Lysine 27 acetylation (H3K27ac; 9E2H10), respectively, do not colocalize with their target epitopes in the nucleus (H2B-Halo; magenta). **B.** Predicted Alphafold3 structure of design 13 of the anti-H3K27ac scFv (9E2H10) bound to the H3 tail with Lysine 27 acetylated (color coded by pLDDT). **C.** The difference in predicted pLDDT scores between Alphafold3 models with an H3 peptide harboring acetylation (as in B) versus those without. The first 44 amino-acids of the H3 tail are shown along the x-axis, with a dashed line marking K27ac. **D.** Image of a cell expressing design 13 of the anti-H3K27ac scFv pre- (top) and post-TSA (bottom). **E.** Left, the nuclear to cytoplasmic ratio of anti-H3K27ac scFv (design 13) in living cells post treatment with the deacetylation inhibitor TSA (purple), the vehicle control DMSO (green), or the p300 inhibitor A485 (orange). Nuclear FRAP recovery curves from cells, as in E (3 hrs post inhibitor treatment). **F.** Predicted Alphafold3 structure of design 4 of the anti-H3S10ph scFv (7G1G7) bound to the H3 tail with Serine 10 phosphorylated (color coded by pLDDT). **G.** The difference in predicted pLDDT scores between Alphafold3 models with an H3 peptide harboring phosphorylation (as in F) versus those without. The first 44 amino-acids of the H3 tail are shown along the x-axis, with a vertical dashed line marking S10ph. **H.** Snapshots from a movie of cells expressing the anti-H3S10ph scFv (design 4) during mitosis. Scale bars, 10 μm.

For each antibody sequence, we generated 15 candidate scFvs with our pipeline and chose the top four or five by pLDDT to screen in living cells. To see if modeling the target epitope was critical, we tested a couple of strategies. For H3K27ac, we inputted the scFv along with the target peptide (the H3 N-terminal tail with glutamine at position 27 to mimic acetylation), while for H3S10ph and H3K9me3, we just input the scFv sequence without the target modified peptide. Although we could have used AlphaFold3^62^ to more properly model post-translational modifications, we chose AlphaFold2 because at the time it was more open and could be used without restriction.

Of the four designed anti-H3K27ac mintbodies we screened, one localized in the nucleus of U2OS cells, where histone acetylation is expected to be (Figs. S3A). Reassuringly, when this sequence was later input into AlphaFold3 together with its target H3 tail peptide (Fig. 3B), the model confidence (pLDDT) of the H3 peptide peaked at K27 when that residue was acetylated, even when competing acetylation marks were nearby (Fig. 3C). To more directly test specificity in an experimental setting, we tracked the cellular localization of the mintbody in response to the addition of the histone deacetylase inhibitor trichostatin A (TSA; to increase H3K27ac) and the p300 histone acetyltransferase inhibitor A485 (to decrease H3K27ac). As expected of a functional anti-H3K27ac intrabody, the ratio of nuclear to cytoplasmic signal increased in the presence of TSA (Fig. 3D and Fig. 3E, purple; Movie S1) and decreased in the presence of A485 (Fig. 3E, orange and Movie S2). As well, FRAP recoveries were faster in TSA than they were in control cells, and faster yet in A485-treated cells, indicating progressively fewer target binding sites (Fig. 3E, right). The FRAP recovery (t_1/2_ ∼ 15 sec) was also on par with similar measurements using purified fragmented antibodies (Fab) physically loaded into cells (t_1/2_ ∼ 21 sec)^63^, indicating the CDRs are largely intact in this mintbody.

Regarding anti-H3S10ph, four out of five scFv designs localized with mitotic chromatin, as would be expected (Fig. S3B). One design (design 4) had the best signal-to-noise. When we input this sequence into AlphaFold3 together with its target H3 tail peptide (Fig. 3F), the model confidence (pLDDT) of the H3 peptide peaked at S10 when that residue was phosphorylated, even when competing phosphorylation marks were nearby (Fig. 3G). This success suggested the epitope may not be necessary in the original model used to generate the mintbody designs. Time-lapse imaging of design 4 in living cells undergoing mitosis revealed the rapid dynamics of H3S10 phosphorylation and dephosphorylation (Fig. 3H). The anti-H3S10ph mintbody marked phosphorylation in patchy spots across the nucleus nearly five minutes before mitosis could be detected based on histone H2B localization (Movie S3), a result also seen using purified Fab physically loaded into cells^60^. Loss of phosphorylation was also rapid as cells approached telophase, with the condensed mitotic chromatin closest to the metaphase plate harboring phosphorylation longest.

Unfortunately, all designs we tested for the anti-H3K9me3 mintbody failed to properly localize within the nucleus (Fig. S3C), despite having high pLDDT scores according to AlphaFold2 and 3. Some designs were worse than others, however. In particular, design 1 could not be detected in cells at all, and design 3 showed characteristic aggregates that are most likely indicative of misfolding. It remained unclear why this particular antibody suffered, although a clue may come from the parental antibody (2F3/CMA318), from which it was difficult to generate Fab from due to the presence of protease susceptible sites in the heavy chain^52^. It is possible these sites may also hinder scFv folding, stability, and overall performance.

### Developing a panel of intrabodies against diverse histone modifications

As a final demonstration of the practicality and speed of our approach, we applied our scFv pipeline to a full panel of histone modification-specific mouse monoclonal antibodies that were previously developed, sequenced, and validated for binding specificity and strength^52,59,61,64–68^. We were especially curious if there are specific histone modifications that are inherently difficult to target. Among the 19 antibody sequences we ran through the pipeline, 14 resulted in successful scFv, having at least one design that exhibited enrichment in nuclei or mitotic chromosomes (Figs. 4A and S4). For example, the original anti-H4K16ac scFv failed to localize to the nucleus, but 4 out of 5 scFv designs exhibited nuclear enrichment (Fig. 4B). Although the nuclear to cytoplasmic ratios were high for designs 2 and 5 (Fig. 4C), their expression levels – both the intensity and the number of expressing cells – were lower than design 3, as revealed by Western blotting (Fig. 4D). Other acetylation-specific antibodies (H3K14ac, H4K5ac, H4K8ac, and H4K12ac) were also vastly improved, exhibiting strong nuclear enrichment that was completely lacking in their original counterparts (Fig. S4A).

**Fig. 4.**
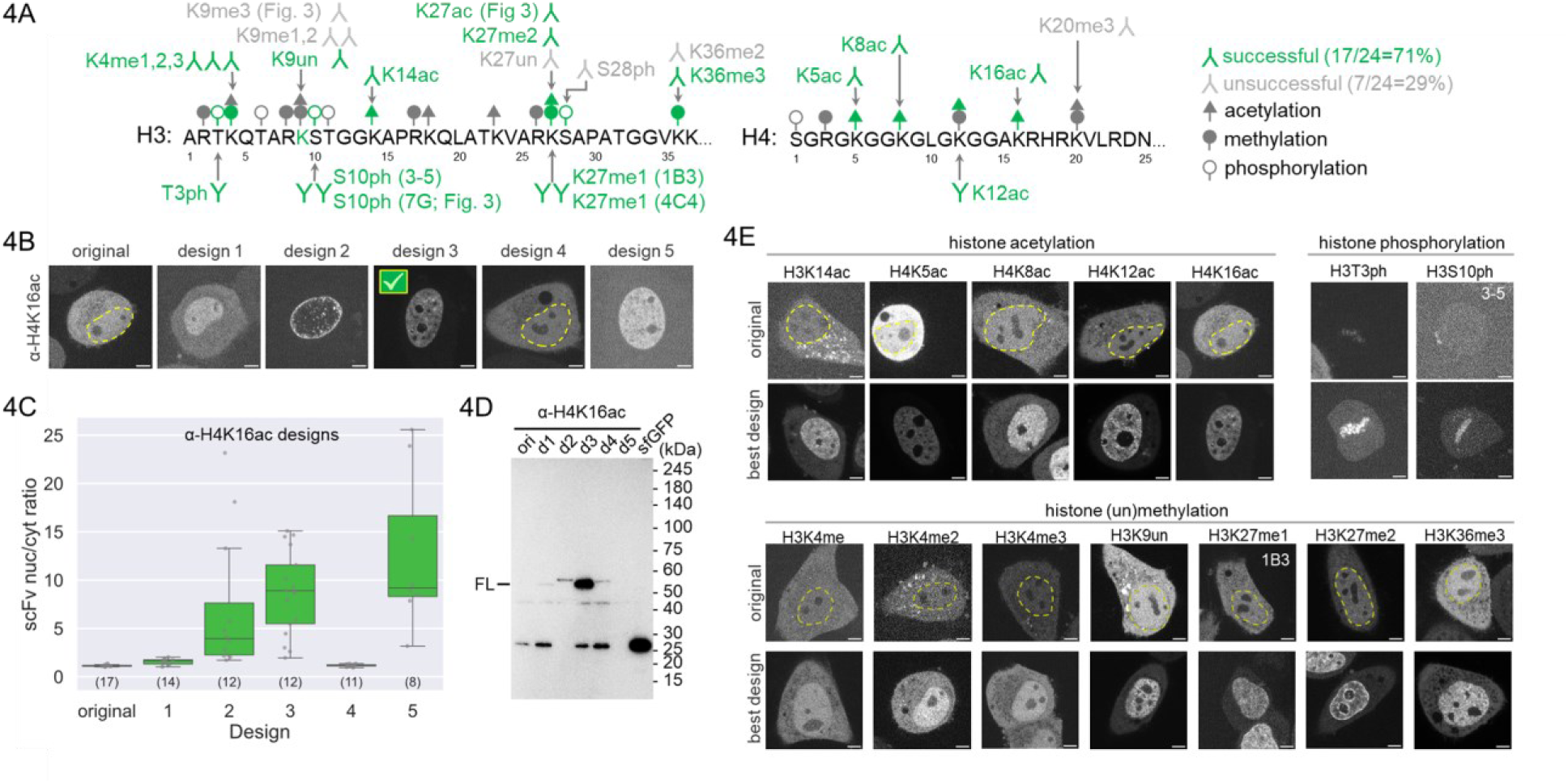
A panel of intrabodies targeting diverse histone modifications. **A.** 24 antibodies against histone H3 and H4 post-translational modifications were passed through the pipeline to generate candidate mintbodies (modification-specific intrabodies). Top-ranked (by pLDDT) candidates (4-5 per antibody) were tested in living HeLa cells. 17 of 24 attempts were successful (highlighted in green versus gray). **B.** Top, sample images of cells expressing candidate mintbodies against H4K16ac to compare which localize better to the nucleus (which is marked by a dashed yellow line when needed). Design 3 showed the best localization pattern. **C.** Sample live-cell measurement of the nuclear to cytoplasmic ratio of the anti-H4K16ac mintbody designs (number of cells (N) indicated; box is 25-75% interquartile range; whiskers mark points within 1.5✕ the interquartile range). **D.** A Western blot showing that design 3 (d3) of the anti-H4K16ac mintbody has the highest full-length (FL) expression level in cells. **E.** Sample live-cell images of each successful mintbody comparing the localization of the original mintbody (top rows) compared to the best design (bottom rows). The best designs localize better to the nucleus (which is marked by a dashed yellow line when needed) or, in the case of phosphorylation, to mitotic chromatin. n.d., mitotic cells not detected. Scale bars, 5 μm.

Among the 12 methylation-specific antibodies we tested, 8 localized to the nucleus, including three targeting monomethylation (H3K4me1 clone 19A5/CMA302 and H3K27me1 clones 4C4 and 1B3/CMA321), two targeting dimethylation (H3K4me2, clone 27A6/CMA303 and H3K27me2 clone 5A12/CMA322), and two targeting trimethylation (H3K4me3 clone 16H10/CMA304 and H3K36me3 clone 2C3/CMA333) (Fig. S4C). This suggests our inability to target H3K9me3 (Fig. S3E) was most likely due to chance rather than being indicative of an inherent problem targeting methylation more generally.

Of the two phospho-specific antibodies we ran through our pipeline, one (anti-H3S10ph clone 3-5/CMA313) clearly marked mitotic chromosomes (Fig. S4B). To further test our pipeline, we also examined an scFv that was functional from the start (anti-H3T3ph clone 16B3). In this case, all five pipeline designs localized to mitotic chromosomes like the original (Fig. S5A). Although these designs did not have superior signal-to-noise compared to the original (Fig. S5B), the fact that all five retained functionality highlights the robustness of our pipeline.

We hypothesized that the success of our designs stemmed from improved solubility and stability. To test this directly in a controlled setting, we expressed anti-H4K16ac mintbodies (1B2/CMA416) using an *in vitro* translation system. After centrifugation, the original mintbody largely precipitated, whereas the redesigned version was more evenly distributed between the supernatant and precipitate (Fig. S5C). Furthermore, nearly all of the original mintbody in the supernatant precipitated after heating to 60°C and 70°C, whereas a substantial fraction of the redesigned version remained soluble under the same conditions (Fig. S5C). These results suggest that scFvs redesigned using our pipeline have greater thermal stability than their original counterparts.

Finally, we generated stable polyclonal cell lines for each of our 17 newly designed mintbodies in human (HeLa or U2OS) and mouse (NIH3T3) cells. The successful establishment of these cell lines indicates that the new mintbodies can be expressed at an appropriate concentration without overtly disrupting cellular viability. Mintbodies bind their targets reversibly and in a monovalent manner, which limits the duration of target engagement compared to full-length antibodies^27^. However, at higher concentrations, they may still affect endogenous interactions, depending on the experimental conditions.

## DISCUSSION

In this study, we used AI-based tools to accelerate the development of functional intrabodies by modeling and optimizing their framework regions. Much effort has been made to evolve and design CDRs to obtain antigen binders with high specificity and affinity. In contrast, although the framework regions play a critical role in securing antibody structure and stability, efforts to develop a general strategy to improve their intracellular function have been limited^14,69^. By focusing on refining these framework regions while leaving the CDRs intact, we achieved a high success rate in converting existing antibody sequences into functional intrabodies.

Out of the 26 parental antibody sequences we tested, 19 were successfully converted into functional scFv intrabodies, yielding an overall success rate of approximately 73%. This success rate is significantly higher than the 5-10% success rate we’ve experienced when directly converting endogenous antibody sequences. Importantly, we began with designs that all initially failed as scFv intrabodies (aside from anti-H3T3ph), many of which also failed after standard loop grafting onto stable scaffolds having high sequence similarity, usually an indicator of grafting success^7^. Despite these difficult starting points, out of a total of 127 designs the pipeline generated, 51 functioned effectively in living cells, so the average probability that any single design worked was roughly 41%.

These statistics highlight the effectiveness of our pipeline to optimize scFv framework regions for intracellular expression. As a first application, we created intrabodies against the popular synthetic FLAG tag and an epitope within the SARS-CoV-2 nucleocapsid protein. We then focused on histone modifications, tripling the number of intrabodies available for tracking the dynamics of these elusive marks *in vivo*, a traditionally challenging task^11,27^. With these intrabody sequences now stored on plasmids, it is easier to distribute them through platforms like Addgene and to modify them, for example, to create bispecific antibodies that could be used to explore the combinatorics of the histone code hypothesis^24,70,71^. These applications will complement existing tools like GFP-tagging, further expanding our ability to probe complex biological processes.

The potential impact of our approach is substantial. With over 2,000 solved antibody structures available in the Structural Antibody Database^72^ (SAbDab) and more than 147,000 antibody sequences from the Observed Antibody Space^73^ (OAS), each represents a candidate for conversion into a functional intrabody. Importantly, our pipeline requires only the antibody sequence, not the structure, and can even work without inputting the target epitope sequence. This is ideal because the number of publicly available antibody sequences is rapidly growing, as researchers prioritize preserving these sequences to address the antibody crisis caused by variability in commercial antibody batches^74,75^. Our method offers a timely approach to take advantage of this expanding resource.

Interestingly, our results suggest that while AI-based tools like AlphaFold offer valuable structural predictions, they alone are insufficient for ensuring successful intrabody design. For example, high pLDDT scores, which indicate predicted structural confidence, did not consistently correlate with functional success in our live-cell screens. This finding highlights the importance of experimental validation and suggests a need for new metrics that better predict intrabody functionality. Notably, the failures in our pipeline were just as informative as the successes. By comparing successful and unsuccessful designs in anti-EpNP scFv, we identified a key residue within the antibody framework that had previously been shown to play a crucial role in stabilizing scFv structure in living cells^55^. In contrast, we observed that some antibodies, such as anti-H3K9me3 or anti-H3S28ph, consistently failed for reasons that remain unclear, whether due to issues with the parental antibody or other hidden factors. Despite these occasional failures, we believe our approach will be instrumental in mapping the key residues that stabilize scFv structures, thereby improving future intrabody designs.

While our work was being prepared, new methods such as AbMPNN^76^, IgMPNN^37^, and AntiFold^35^ have emerged, building on ProteinMPNN^42^ and AlphaFold2^29^ to improve antibody structure prediction and design by incorporating antibody-specific data. Although these tools focus on optimizing CDRs to improve binding affinity^37,76^ and also prevent self-binding and non-specific targeting^38^, the extent to which these tools can also improve the framework regions of intrabodies for functionality inside living cells remains unclear. Developing tools to optimize these regions will be useful, and comparing their predictions with our experimentally validated designs will offer insights into what drives intrabody success in living cells.

To facilitate these sorts of comparisons, we provide a comprehensive list of all sequences we designed and tested, ranked by visualization and quantification in living cells (Table S1). As more designs are generated and analyzed, this resource will grow, making it easier to distinguish successful from unsuccessful intrabody designs. Ultimately, we hope to train improved algorithms with this data, with the goal of developing a next-generation pipeline specifically trained on functional intrabodies to further reduce the need for labor-intensive live-cell screening.

## MATERIALS AND METHODS

### Antibody to Intrabody Pipeline

Our pipeline to convert antibody sequences into designed-intrabody sequences consists of five steps. First, we aligned and annotated the antibody heavy and light chain sequences, typically using Clusta^l77^ for alignment and ANARCI^40^ or Benchling’s built-in antibody property prediction tool for CDR and framework region annotation. We sometimes compared predicted CDRs and chose a conservative set of residues encompassing all predictions. Second, based on the annotations from step one, we designed the scFv sequence, starting with the variable region of the heavy chain at the N-terminus (containing four framework regions and three CDRs), followed by a GS-rich linker and ending in the variable region of the light chain (also containing four framework regions and three CDRs). Third, a structure was predicted for this sequence with AlphaFold2^29^ using ColabFold^41^. Fourth, this structure was used as input into ProteinMPNN^42^. During the ProteinMPNN sequence design the fixed positions included the CDRs as well as all residues in the scFv’s framework regions that were within 3Å of the CDRs, along with the GS-rich linker. We experimented with fewer angstroms when designing M2 anti-FLAG scFvs, but this seemed to lower the success rate (e.g. we tested 0 Å with the M2 anti-FLAG antibody and this produced 0 successful intrabodies). By contrast, a value of 5Å left important residues we had previously identified in the framework regions unchanged (e.g., M83, which we previously discovered^47^ was an indicator of non-functional scFv). The remaining residues form the framework of the scFv, to be redesigned by ProteinMPNN. The settings were: Number of sequences = 15, Sampling temperature = 0.1, Model = vanilla–v_48_020 or soluble–v_48_020, Backbone noise = 0.02. Fifth, we predicted the structure of all ProteinMPNN guided sequence designs using ColabFold and ranked these by pLDDT.

We provide python code on Github that implements our complete pipeline in python (https://github.com/jbderoo/scFv_Pmpnn_AF2), using ANARCI^40^ to annotate CDRs (using the Martin numbering scheme by default), LocalColabFold^41^ to predict intrabody structures, and ProteinMPNN^42^ to generate new framework sequences for each input.

### Plasmid Construction

The nucleotide sequence of histone modification-specific antibodies was determined as previously described^78^. AI-assisted scFv library designs were reverse translated into human codon optimized DNA sequences. The library scFv DNA sequences were flanked with sequences homologous to the PB533_15F11-sfGFP vector (Addgene Plasmid #167526) to be inserted at the NheI/AgeI cut sites with isothermal assembly. scFv sequences were synthesized by TwistBioscience. In some cases, scFv sequences were synthesized by IDT (eBlocks) and cloned into the PB533 vector (System Biosciences) harboring an sfGFP sequence^57^ using the In-Fusion Cloning system (Takara Bio). Plasmids for all successful scFvs will be made available at Addgene upon acceptance of the manuscript.

### Stable cell line generation

Stable cell lines were generated using the PiggyBac transposon system. All scFv vectors contain PiggyBac LTRs with neomycin resistance for selection. Cells were co-transfected with Super PiggyBac transposase (NovoPro or Systems Biosciences) at 1:5 transposase to transposon in a 35-mm dish at 50% confluency. Once cells reached confluency, cells were expanded into 10-cm dishes and selected for a week with G418 750 μg/mL (or 1 mg/mL for HeLa). Individual colonies were hand picked for monoclonal cell lines or pooled together for polyclonal lines.

### Cell Culture

Human U2OS osteosarcoma or HeLa cells were grown in DMEM (ThermoFisher Scientific or Nacalai Tesque) supplemented with 10% v/v FBS (Atlas or ThermoFisher Scientific), 1% v/v Pen/Strep (ThermoFisher Scientific or Sigma), and 1 mM L-glutamine (ThermoFisher Scientific or Sigma) at 37°C in 5% CO2. Mouse fibroblast NIH3T3 cells were grown in DMEM (ATCC 30–2002) supplemented with 10% v/v FBS (EqualFETAL, Atlas Biologicals) and 1% v/v Pen/Strep (Gibco 15140122) at 37°C in 5% CO2. Trypsinization for cell passaging was performed using TrypLE Express (Thermo Fisher 12605010).

### Transfection

The day prior to imaging, cells were seeded on 35-mm MatTek Chambers or 24-well glass-bottom plates (AGC Techno Glass) at ∼70% confluency and transfected. Transfection was performed using Lipofectamine LTX or 2000 (Thermo Fisher Scientific) with 0.8-1 μg total plasmid DNA. Cells were washed 3 times with DMEM the next morning. In experiments using HaloTag, staining was performed using 200 nM of JF646-HaloLigand for 30 min then washed 3 times with DMEM. Cells were then kept in supplemented DMEM lacking phenol red or FluoroBrite (ThermoFisher Scientific) containing 10% FBS for imaging.

For colocalization imaging experiments (Figs. 1B, 1E, S1A, 2A, 2D, and S2A), the cells were seeded on 35-mm MatTek chambers with 70% confluency. The next day, 1.25 μg of the scFv-GFP construct and 1.25 μg of the epitope-tagged mCh-H2B construct were transfected into cells using Lipofectamine LTX. The medium was changed to complete DMEM medium at 3 hours post transfection. The cells were imaged at 20 h post transfection. Cell treatment for anti-H3K27ac design 13 imaging experiments were performed as above except 2.0 μg of scFv-GFP and 1.0 μg of mCh-H2B constructs were used (Figs. 3D, 3E).

### Western blotting

For Western blotting analysis in Fig. 4C, cells were plated on 12-well plates (Corning, 3513) at 1.0 x 10^5^ cells/well and transfected with 1.6 μg plasmid DNA using Lipofectamine 2000 (Thermo Fisher Scientific). The next day after transfection, cells were washed 3 times with phosphate-buffered saline (PBS) and lysed with 1X SDS-gel loading buffer (2% SDS, 10% glycerol, 100 mM Tris-HCl [pH 6.8], 100 mM dithiothreitol, 0.01% bromophenol blue; 150 μL/well), before incubating at 95°C for 5 min.

For Fig. S5B, mintbodies were produced by an in vitro transcription-translation coupled system using PURE frex 2.0 (Gene Frontier). Approximately 90 ng of each template DNA in a 20 μL reaction mixture was incubated for 6 h at 37°C. After centrifugation at 16,000 ×*g* for 30 min at 4°C, supernatant was transferred to a new tube. After incubating supernatant fractions for 30 min at 50, 60 and 70°C, precipitated materials were removed by centrifugation as above. All fractions were added with an equal volume of 2X SDSample buffer and heated for 10 min at 95 °C.

Protein samples were separated in polyacrylamide gels (SuperSep Ace, 5-20 %, 17 well; Fujifilm Wako) and transferred onto a 90 mm x 90 mm polyvinylidene difuoride membranes (FluoroTrans W; Fujifilm Wako) using a semi-dry system (Trans-Blot Turbo; Bio-Rad; 1.3 A for 15 min) with EzFastBlot (Atto) as a transfer buffer. Membranes were washed with TBST (20 mM Tris-HCl [pH 8.0], 150 mM NaCl, and 0.05% Tween 20), blocked for 20 min in Blocking-One (Nacalai Tesque) with gentle shaking, and washed with TBST. Membranes were then incubated with mouse anti-GFP antibody (GF200, Nacalai Tesque; 1:2,000 dilution) in Can-Get-Signal Solution 1 (Toyobo) for 1 h at room temperature, washed three times with TBST for 5 min each, incubated with horseradish peroxidase-conjugated sheep anti-mouse Ig (Jackson ImmunoResearch; 1:10,000 dilution) in the Can-Get-Signal Solution 2 (Toyobo) for 1h at room temperature, and washed three times with TBST for 5 min each. Chemiluminescence signals were developed using ImmunoStar LD (Fujifilm Wako) and detected using a luminescent imager LuminoGraph II (Atto).

### Live-cell Imaging

#### Olympus Spinning Disk Confocal Microscope

For Figs. 1B,E-G, S1A (top), 2A,D-F, S2A, 3D-E, S3A (right), and Movies S2, images were captured on an Oympus IX8 Spinning Disk Confocal Microscope (CSU-22 head) equipped with an iXon Ultra 888 EMCCD camera (Andor) operated by SlideBook (Intelligent Imaging Innovations) coupled with a Phasor photomanipulation unit (Intelligent Imaging Innovations), and a 60x (NA 1.42) oil immersion objective. Cells were enclosed in a custom-built 37°C chamber with 5% CO_2_ throughout imaging. A 488-nm laser was used to capture GFP tagged scFv’s and a 561-nm laser was used to capture H2B-mCh.

Imaging for 1F and 2E were acquired at full cell volumes with 17 z-stacks at 0.5-μm step size with the 488-nm laser set to 50% power and 100 ms and the 561-nm laser at 30% power with 100 ms exposure. Time lapse imaging in 3E (left) was captured with 21 z-stacks at 0.5-μm step size and 5 min interval for 175-200 min. The 488-nm laser was set to 40% power at 200-ms exposure time, and the 561 nm laser was set to 20% power at 100 ms exposure time. FRAP experiments (1G, 2F, and 3E right) were performed on cells transiently transfected (as described above). Single plane imaging was performed about the center of the nucleus. Before photobleaching, 10 frames were acquired at 1s time interval. Between frames 10 and 11 the Phaser 488-nm laser was set to bleach a circle of ∼1 μm in diameter for 100 ms at 90% power. Image captures continued every 1s for an additional 170-250 times points.

For drug treatment imaging and FRAP of U2OS cells expressing anti-H3K27ac mintbody (Fig. 3D and 3E), the position of cells expressing both the scFv and mCh-H2b were recorded and an initial image was taken. Cells were imaged in 2.5 ml of media. 0.5 ml of media was removed and mixed with 2 μl of drug in DMSO or DMSO only, and then added back to the imaging dish. Working:Final concentrations of TSA and A485 was 625 μM:500 nM and 12.5 mM:10 μM respectively. Starting 5 minutes after drug addition, the cells were imaged every 5 minutes for 2 to 3 hrs. Cells which were not imaged during the 3 hr time lapses were used for the FRAP experiments as described in the previous section.

#### Custom HILO microscope

For S1A (bottom) and S3A imaging, a custom built high inclined thin illumination fluorescence microscope^79^ equipped with a 60x (NA 1.49, Olympus) oil-immersion objective lens, three solid-state laser lines (488, 561, and 637 nm, Voltran), a T660lpxr ultra-flat imaging grade emission image splitter (Chroma), and two iXon Ultra 888 EMCCD cameras (Andor) was used. A 300 mm achromatic lens (Thorlabs) was used to focus images onto the camera instead of the standard 180 mm Olympus tube lens, resulting in an effective magnification of 100x. Cells were incubated in a 37°C chamber with 5% CO_2_ (Okolab). Images were captured at 512 x 512 px^2^ size, 53.64 msec exposure time. A single z plane was captured about the center of the nucleus.

#### Dragonfly and Yokogawa CSU-W1 Microscopes

For Fig. 3A and S3-5, a spinning-disk confocal system (Andor DragonFly or Yokogawa CSU-W1 for Fig. S3B) with an inverted microscope (Ti-E; Nikon), operated by Fusion software ver. 2.2.0.50 (Andor) or NIS-elements software ver. 5.11.03 (Nikon; for Fig. 3H and S3B), was used with a heated stage at 37°C supplemented with 5% CO2 (Tokai Hit). Single optical sections were acquired with a 100x PlanApo VC (NA 1.4) oil-immersion objective lens, a 405/488/561/640 or a 405/470/555/640NIR (for Fig. S3B) dichroic mirror, a 521/38 emission filter or a 520/60 emission filter (for Fig. S3B), a laser unit (5-20% transmission of 488 nm; LC-ILE-400-M; Andor or 30% transmission of 470 nm with a 10% neutral density filter; LDI-7 Laser Diode Illuminator; Chroma Technologies Japan for Fig. S3B), and an EM-CCD (iXonUltra; Andor; gain multiplier 300; exposure time 0.3-0.5 s or iXon+; Andor; gain multiplier 300; exposure time 0.1-1 s for Fig. S3B). Transmission images were also acquired.

For time lapse imaging (Fig. 3H and Movie S3), HeLa cells stably expressing sfGFP-tagged anti-H3S10ph (7G1G7) scFv (design 4) and H2B-Halo^55^ were plated on a 35-mm glass-bottom dish (AGC Technology Solutions). The next day, the cells were stained with 100 nM Janelia Fluor 646 HaloTag Ligand (Promega) for 1 h, before replacing the medium with FluoroBrite containing FBS (Thermo Fisher Scientific) and setting onto a heated stage at 37°C supplemented with 5% CO2 (Tokai Hit) featured on a spinning-disk confocal system (Yokogawa CSU-W1) with an inverted microscope (Ti-E; Nikon), operated by NIS-elements software ver. 5.11.03 (Nikon). Images were acquired with a 100x PlanApo VC (NA 1.4) oil-immersion objective lens, a 405/470/555/640NIR dichroic mirror, 520/60 and 690/50 emission filters, a laser unit (20% transmission of 470 nm and 15% of 640 nm with a 10% neutral density filter; LDI-7 Laser Diode Illuminator; Chroma Technologies Japan), and an EM-CCD (iXon+; Andor; gain multiplier 300; exposure time 500 ms). Fifteen different focal planes were imaged at 1 μm intervals every 5 min. Max intensity projection images are shown in Fig. 3H and Movie S3.

### Image Quantification

#### Nuclear/Chromatin to cytoplasmic ratio

The ratio of nuclear to cytoplasmic signal intensity of cells from movies (Figs. 1F, 2E, and 3E, left) was calculated using custom scripts (Nuc2Cyt scripts on (https://github.com/Colorado-State-University-Stasevich-Lab/StasLabImageAnalysis). The maximum intensity projection of image stacks of full cell volumes was used for quantification. Background fluorescence was removed by subtracting the 1st percentile of intensity values, and negative values were set to zero. Nuclei were identified using Otsu thresholding of H2B and a binary mask was created to measure the nuclear signal. To measure cytoplasmic signals, a ring mask was generated around each nucleus with a 6-pixel width. The average intensity of intrabodies for both nuclear and cytoplasmic regions was measured and the ratios were then calculated. For live cell tracking experiments (Fig 3D, 3E Left), each cell in a time lapse was given a unique identifier using Trackpy which was used to link the nuclear and cytoplasmic signals of individual cells for the duration of the experiment.

For Fig. 4C, and S3-5, nuclear and cytoplasmic areas in mintbody-expressing cells were selected manually, assisted with transmission images of cells. After the background intensity in a cell-free area was subtracted from the nuclear and cytoplasmic intensities, the nuclear to cytoplasm intensity ratio was determined.

#### Fluorescence Recovery After Photobleaching (FRAP)

FRAP curves (Figs. 1G and 2F) were calculated using a custom script written in Python (https://github.com/Colorado-State-University-Stasevich-Lab/frap_analysis). Briefly, the FRAP spot location for the measurement was determined by subtracting the first post-FRAP frame (frame 11) from the previous frame (frame 10). The resulting image was used to create a mask that marked the bleached area, and a Difference-of-Gaussians filter was applied to further enhance the precise location of the center of the FRAP spot. A circular region was then inscribed into the bleached area, and the average signal from this region was plotted for the duration of the experiment (normalized to the intensity from frames 1-10).

## ACKNOWLEDGEMENTS

We thank Prakash Shyam Karuppiah and Jacob Leavitt for assistance with preliminary experiments during their rotation in the Stasevich lab. We thank the Jennifer Deluca lab (Colorado State University) for helpful scientific discussions. We thank Harumi Ueno (Tokyo Institute of Technology) for technical assistance and the Center for Integrative Biosciences and the Biomaterials Analysis Division, Open Facility Center at Tokyo Institute of Technology for DNA sequencing. We note that ChatGPT was used to assist with text grammar and clarity.

