## Supplementary Figures and Movie Legends for "AI-assisted protein design to rapidly convert antibody sequences to intrabodies targeting diverse peptides and histone modifications"

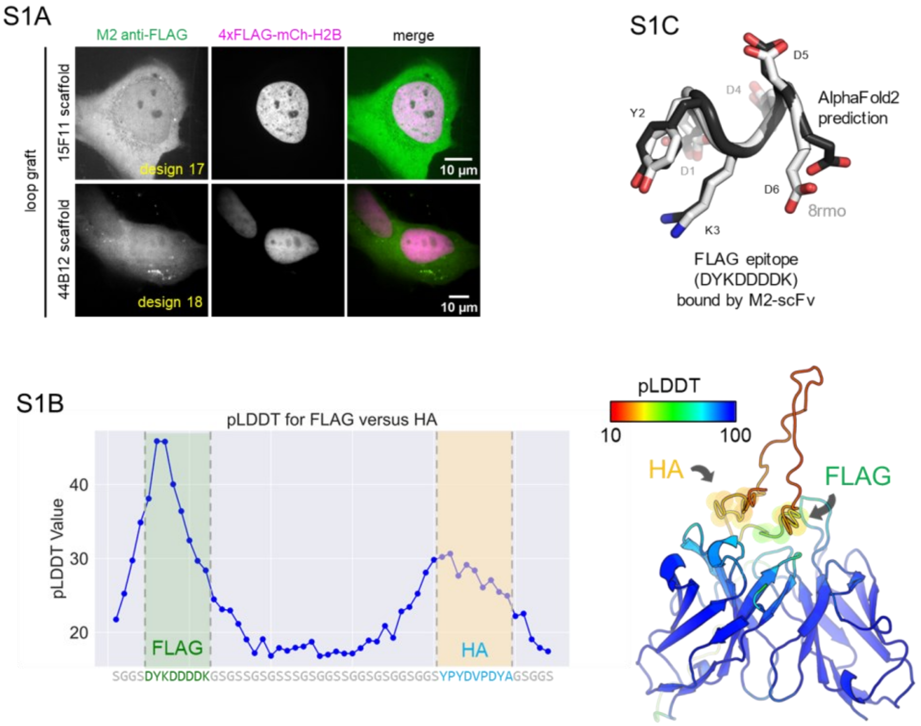


**Fig S1. Testing anti-FLAG scFv designs. A.** Loop-grafted (top: 15F11 scaffold; bottom 44B12 scaffold) M2 anti-FLAG scFv (green) do not colocalize with their target FLAG epitope (4xFLAG-mCh-H2B; magenta) in live cells. **B.** The Alphafold2 predicted pLDDT scores of an unstructured binder harboring an HA and FLAG epitope when modeled with the original M2 anti-FLAG scFv (design 0). **C.** An overlay of the AlphaFold2 predicted structure of the FLAG epitope when bound by the M2 anti-FLAG scFv (design 12) compared to the experimentally determined structure of the FLAG epitope when bound by the M2 anti-FLAG Fab fragment (8RMO).

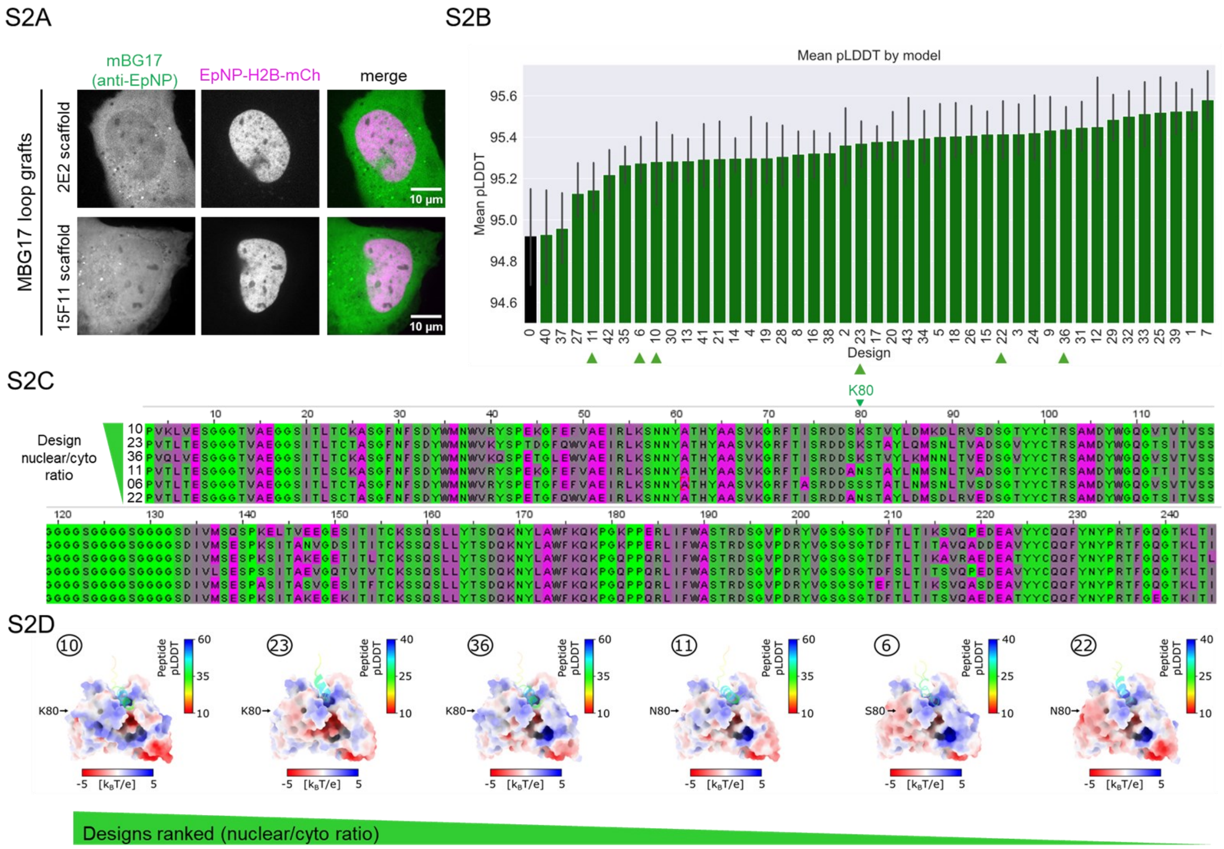


**Fig. S2. Testing an scFv against an epitope in the SARS-CoV-2 nucleocapsid protein. A.** Loop-grafted (top: 2E2 scaffold; bottom 15F11 scaffold) scFv (mBG17; green) against the SARS-CoV-2 nucleocapsid protein do not colocalize with their target epitope (EpNP; EpNP-mCh-H2B; magenta) in live cells. **B.** The ranked mean pLDDT scores of all mBG17 pipeline designs (includes both scFv and EpNP; green bars) along with the original (black bar). Error bars show the standard deviation from LocalColabFold predicted models (N=5 each). Green arrows indicate designs that were chosen for screening in live cells. **C.** Aligned sequences of six designs for the mBG17 scFv, ranked from top to bottom by the nuclear to cytoplasmic ratio they displayed in living cells, colored by helix propensity. Residue K80 is denoted by a green arrow. **D.** All 6 models are depicted with electrostatic potential at the surface calculated via APBS with default settings in PyMOL. The model used comes from a predicted complex with the peptide partner, but the peptide is excluded from the electrostatics calculation. In all cases, the peptide is placed among the CDR loops despite poor confidence metrics. As shown, the site of residue 80 is distant from the binding cleft and represents one surface change among many across the design panel.

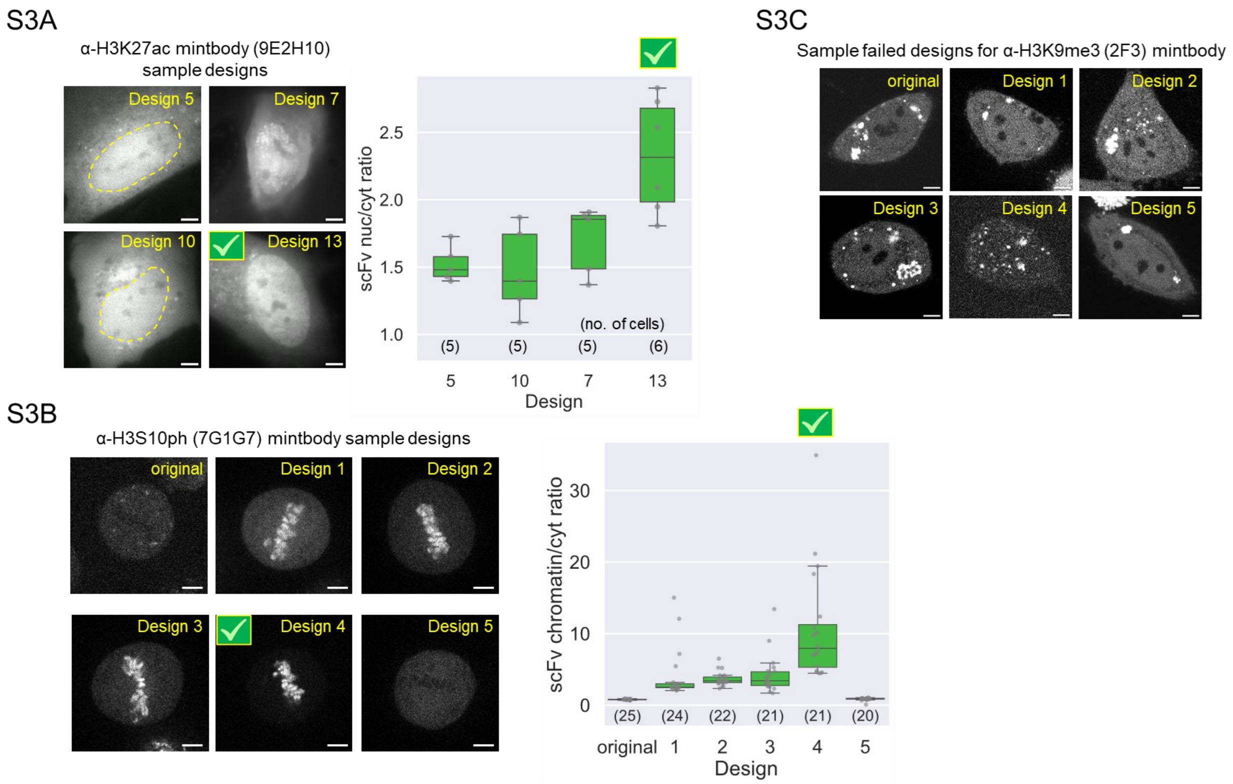


**Fig. S3. Sample images and quantification of H3-specific scFv designs. A.** Sample images of four different designs of an anti-H3K27ac mintbody (based on clone 9E2H10/CMA309). Quantification of the nuclear-to-cytoplasmic ratio for each design is shown on the right, indicating design 13 is best (number of cells (N) indicated; box is 25-75% interquartile range; whiskers mark points within 1.5× the interquartile range) **B**. Sample images of the original and five different designs of an anti-H3S10 mintbody (based on clone 7G1G7/CMA311). Quantification of the mitotic chromosome to cell ratio on the right demonstrates design 7G-4 is best (number of cells (N) indicated; box is 25-75% interquartile range; whiskers mark points within 1.5× the interquartile range). **C.** Sample images of the original and five different designs of an anti-H3K9me3 mintbody (based on clone 2F3/CMA318). The presence of bright aggregates and a lack of nuclear colocalization in all cases indicates the new mintbody designs failed to label target H3K9me3. Scale bars, 5 μm.

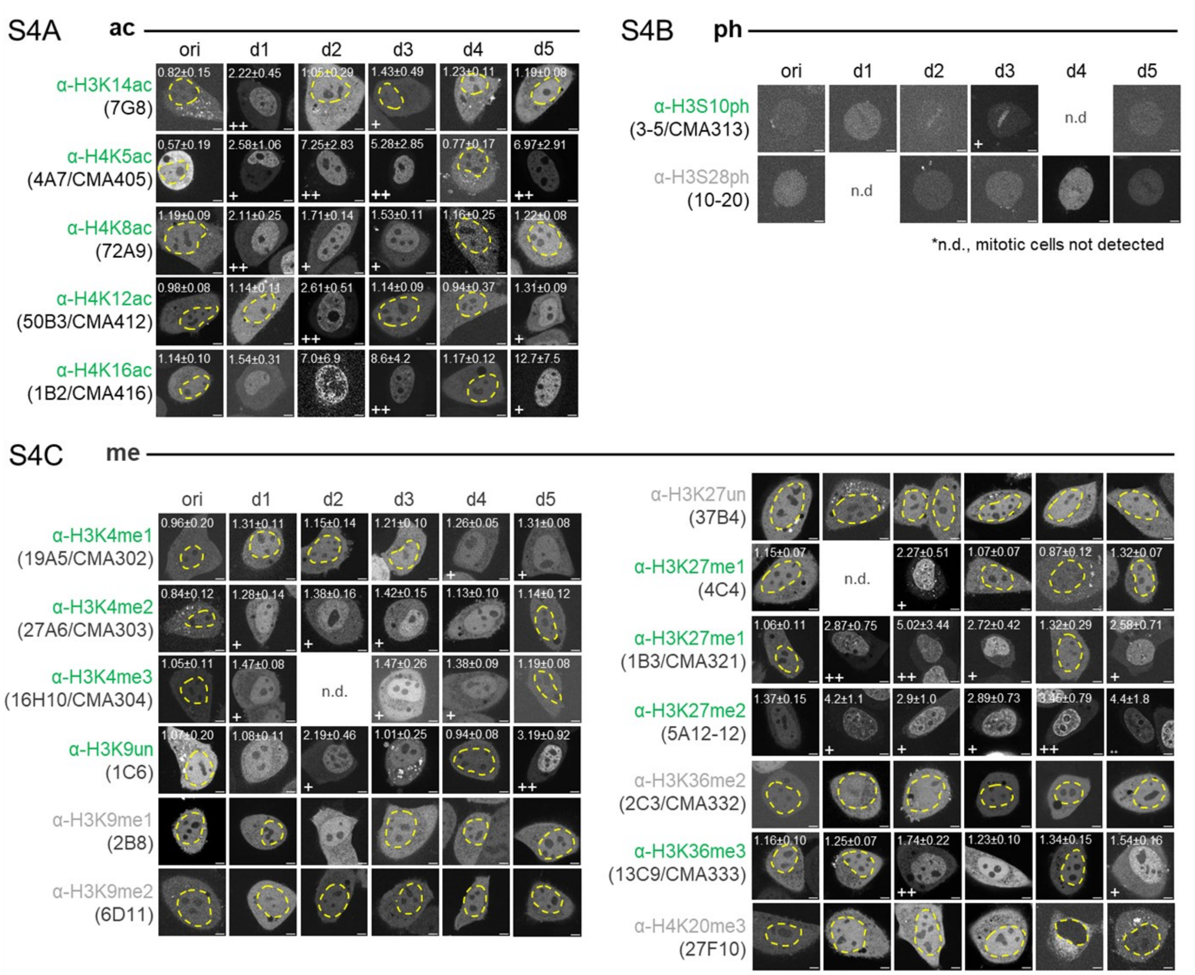


**Fig. S4.** Sample images of live HeLa cells expressing candidate mintbodies (scFv) to compare which localize better to the nucleus (which is marked by a dashed yellow line when needed). **A.** Mintbodies against histone acetylation. **B.** Mintbodies against histone phosphorylation, where localization to mitotic chromatin is compared. **C.** Mintbodies against histone (un)methylation. For successful acetylation and methylation designs, the nuclear to cytoplasmic intensity ratio was measured in living cells and is reported in the upper left of each panel (mean ± s.d.). Some images are reproductions of those in Fig. 4. Scale bars, 5 μm.

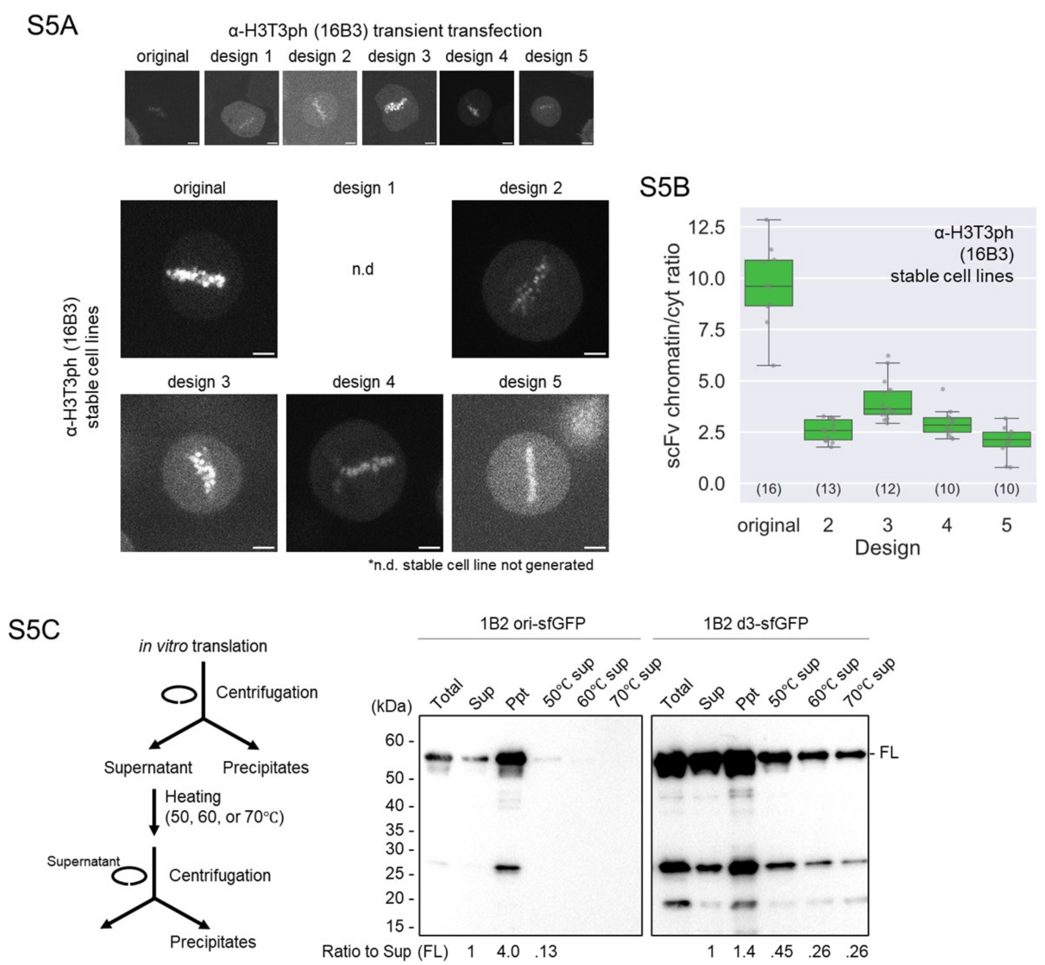


**Fig. S5. Further tests of the robustness of the antibody-to-intrabody pipeline. A.** Sample images of cells expressing candidate mintbodies against H3T3ph compared to the original (top row, transient transfections in HeLa cells; bottom, HeLa cell lines stably expressing each mintbody). In this case the original was functional, and all designs that passed through the pipeline retained their ability to localize to mitotic chromatin (for design 1 we did not generate a stable cell line). **B.** Quantification of the mitotic chromatin to cytoplasmic intensity ratio for data like in A. **C.** Left, flow chart of experiment. Right, Western blot of an *in-vitro* translated anti-H4K16ac mintbody (design 3 of 1B2). FL marks full length protein. Whereas the original (1B2 ori-sfGFP) is predominantly found in the pellet (Ppt) and is lost at higher temperatures, design 3 (1B2 d3-sfGFP) has a large fraction in the supernatant (Sup), and is not lost at high temperatures. Scale bars, 5 μm.

**SUPPLEMENTARY MOVIE LEGENDS:**

**Movie S1: TSA treatment of U2OS cells expressing anti-H3K27ac mintbody (9E2H10 design 13).** Movie corresponding to Fig. 3D, E (left). 500 nM TSA was added just prior to the first movie frame at time = 0 min.

**Movie S2: A485 treatment of U2OS cells expressing anti-H3K27ac mintbody (9E2H10 design 13).** Movie corresponding to Fig. 3E (left). 10 μM A485 was added just prior to the first movie frame at time = 0 min.

**Movie S3: HeLa cells expressing anti-H3S10ph mintbody (7G1G7 design 4) undergoing mitosis.** Movie corresponding to Fig. 3H. Field of view is 75.47 x 75.47 μm.
